# Embedding biomimetic vascular networks via coaxial sacrificial writing into functional tissue

**DOI:** 10.1101/2024.01.27.577581

**Authors:** Paul P. Stankey, Katharina T. Kroll, Alexander J. Ainscough, Daniel S. Reynolds, Alexander Elamine, Ben T. Fichtenkort, Sebastien G.M. Uzel, Jennifer A. Lewis

## Abstract

Printing human tissue constructs replete with biomimetic vascular networks is of growing interest for tissue and organ engineering. While it is now possible to embed perfusable channels within acellular and densely cellular matrices, they lack either the branching or multilayer architecture of native vessels. Here, we report a generalizable method for printing hierarchical branching vascular networks within soft and living matrices. We embed biomimetic vessels into granular hydrogel matrices via coaxial embedded printing (co-EMB3DP) as well as into bulk cardiac tissues via coaxial sacrificial writing into functional tissues (co-SWIFT). Each method relies on an extended core-shell printhead that promote facile interconnections between printed branching vessels. Though careful optimization of multiple core-shell inks and matrices, we show that embedded biomimetic vessels can be coaxially printed, which possess a smooth muscle cell-laden shell that surrounds perfusable lumens. Upon seeding these vessels with a confluent layer of endothelial cells, they exhibit good barrier function. As a final demonstration, we construct biomimetic vascularized cardiac tissues composed of a densely cellular matrix of cardiac spheroids derived from human induced pluripotent stem cells. Importantly, these co-SWIFT cardiac tissues mature under perfusion, beat synchronously, and exhibit a cardio-effective drug response in vitro. This advance opens new avenues for the scalable biomanufacturing of organ-specific tissues for drug testing, disease modeling, and therapeutic use.

## 1. Introduction

Biomanufacturing organ-specific human tissues replete with a biomimetic vascular network remains a formidable challenge.^1^ Without the ability to embed immediately addressable and perfusable vasculature, these engineered human tissues do not remain viable over the time required to provide therapeutic benefit.^[2–7]^ Recent advances in extrusion^[8–10]^, embedded^[3,11–14]^, and light-based^[15–17]^ bioprinting have begun to address this critical need. Yet no method currently allows the free-form patterning of hierarchical, branching vasculature composed of smooth muscle cell-laden shells that surrounds endothelialized lumens in acellular or densely cellular tissue matrices.

Native blood vessels are composed of concentrically arranged layers, in which the inner most layer (intima) is formed by a confluent endothelium that regulates barrier function.^18^ The endothelium is supported by smooth muscle cells (SMCs) which reside in the medial layer and improve vessel robustness.^19^ Coaxial printing is an emerging method for vascular manufacturing, enabling one to directly pattern a sacrificial core (vessel lumen) along with one or more shell materials into concentric layers (vessel wall).^[20–25]^ To date, both bi-layered and tri-layered vessels have been coaxially bioprinted onto a substrate or embedded into acellular support matrices.^[26–29]^ However, these vascular conduits lack the hierarchical branching architectures found in vivo.^[30–32]^

To vascularize organ-specific tissues, Lewis et al. recently developed a method termed sacrificial writing into functional tissue (SWIFT) in which a sacrificial gelatin ink is printed within a granular matrix composed of organ building blocks (OBBs).^3^ These granular OBBs consist of human induced pluripotent stem cells (hiPSCs) in the form of embryoid bodies or differentiated into multicellular spheroids or organoids.^3^ Central to this method is the ability to pattern a sacrificial gelatin ink within these living tissue matrices at low temperature (2 °C), which then liquifies and is washed away upon warming the construct to 37 °C, leaving behind empty channels through which oxygenated culture medium is perfused.^3^ While SWIFT enables fabrication of bulk organ-specific tissues with high cell densities (>10^8^ cells mL^-1^), the embedded perfusable channels lack both the requisite smooth muscle cell-laden wall and confluent endothelial lining of native blood vessels.

Here, we report a generalizable method that unites coaxial embedded and SWIFT bioprinting to pattern hierarchically branching biomimetic vascular networks in both acellular and densely cellular matrices. We first designed a novel coaxial printhead composed of an extended core-shell nozzle. This advance is central to our ability to print hierarchically branching, vascular networks as the extended core feature facilitates its puncture through the shell layer to enable connections to the cores of other printed vessels. Second, we developed a series of core and shell inks to delineate the requisite rheological properties for co-EMB3DP and co-SWIFT printing. To facilitate direct visualization, we carried out initial optimization experiments within a transparent matrix composed of granular alginate microparticles. Next, we printed 3D biomimetic vascular networks via co-EMB3DP in a recently developed microporogen-structured (µPOROS) collagen matrix.^33^ We show that co-EMB3DP ensures seamless integration of sacrificial core and SMC-laden shell features to form 3D hierarchical vascular networks. After printing, the sacrificial core is removed, and the luminal surfaces of these branching vessels are seeded with a confluent monolayer of endothelial cells that provide good barrier function. Finally, after optimizing this method in acellular matrices, we embedded biomimetic vasculature into functional tissue matrices composed of densely cellular, cardiac OBBs derived from hiPSCs via co-SWIFT. Our coaxial bioprinting methods open new avenues for creating vascularized human tissues for drug testing, disease modeling, and therapeutic use.

## 2. Results & Discussion

### Coaxial printhead with extended core-shell nozzle

We first designed a novel coaxial printhead composed of an extended core-shell nozzle with two independently controllable fluidic pathways: for core and shell inks (Figure 1a). This customized coaxial printhead is designed in Solidworks and built using a digital light projection lithography printing. The use of long needles ensures minimal disruption when printing biomimetic vascular networks deep within acellular and densely cellular matrices. The shell ink first travels into an equilibration chamber, which provides a uniform pressure to facilitate ink flow through the shell nozzle. To ensure the shell layer is extruded uniformly, the height and diameter of the equilibration chamber are designed to be roughly one order of magnitude larger (2 mm) than the thickness of this shell layer (0.16 mm). If this criterion is not met, the ink will preferentially flow on one side of the shell layer.

**Figure 1.**
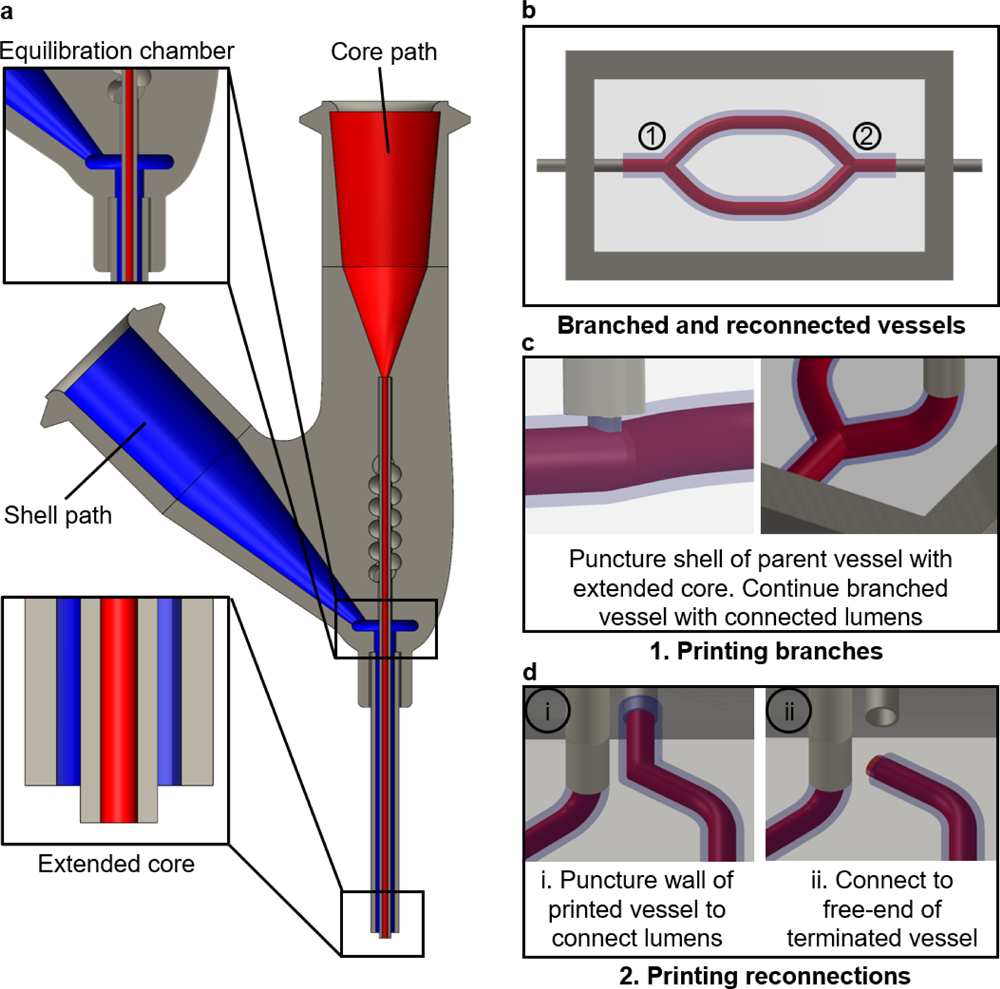
Coaxial embedded 3D printing of branching vessel networks. a) Cross-section of the extended core-shell (coaxial) printhead. b) Schematic of biomimetic vessels composed of bifurcating features embedded within a matrix. c) Schematic illustration connected vessel branches printed using an extended core-shell printhead. d) Schematic illustration of two modes for reconnecting printed vessels.

Creating hierarchically branching core-shell networks requires one to both branch from and reconnect to an existing filament. The core nozzle is extended 250 µm beyond the shell nozzle (roughly the thickness of the shell layer) which is vital for ensuring that the coaxial nozzle punctures the shell layer to connect the cores of the parent and daughter vessels when performing branching and reconnection maneuvers (Figure 1b-d). To create a branch point, one uses the extended core to puncture the shell wall of the filamentary features (Figure 1c). The core and shell inks are then coaxially extruded to ensure connections between the parent and daughter filaments. There are two methods to reconnect printed coaxial filaments. First, the extended core nozzle is once again used to puncture the shell of printed filaments as the core and shell inks are being extruded (Figure 1d,i). Second, a connection could also be formed by connecting at the free end of the printed coaxial filament in a similar manner (Figure 1d,ii). These coaxial interconnections are performed sequentially to build increasingly complex branching vascular networks.

### Optimizing core, shell, and matrix rheology

We created multiple core, shell, and matrix materials to determine the requisite material properties for co-EMB3D printing. We first produced granular alginate particles via an in-air fluidic assembly method (Figure S1). Next, we produced three transparent matrices to enable direct visualization of the co-EMB3DP process and assessed their rheological properties and 3D structure by confocal imaging (Figure S2, Movies S1-S3). Each matrix is composed of granular particles of varying alginate concentration (0.5-2% alginate) and total particle volume fraction (ϕ = 0.80-0.86). We identified the granular alginate matrix (0.5% alginate and ϕ = 0.86) with an intermediate shear yield stress (τ_y_) of ∼70 Pa and the desired viscoplastic and self-healing behavior to be optimal for co-EMB3DP. We then formulated our sacrificial gelatin core ink such that its shear thinning behavior and τ_y_ ∼ 50 Pa nearly matched that of the granular alginate matrix (Figure 2a).^34^ Inspired by the native vessel wall, which is predominantly composed of collagen, we created three shell inks using high-density collagen blended with either gelatin or PBS with τ_y_ values roughly an order of magnitude greater (τ_y_ ∼ 750 Pa), matched (τ_y_ ∼ 40 Pa), or an order of magnitude lower (τ_y_ ∼ 5 Pa) than the alginate matrix. We carried out COMSOL simulations of each shell ink flowing through our coaxial printheads (Figure S3). Each core-shell ink combination could be successfully printed in the vertical direction within this matrix (Figure 2b). However, only the core-shell ink combination with the highest shell τ_y_ ∼ 750 Pa exhibited a uniform core-shell architecture when coaxially printed in the horizontal direction. When the τ_y_ of the shell ink matches that of the matrix, the shell layer thins around the bottom of the filament. When its τ_y_ is less than the matrix, the shell does not entirely wrap around the core ink as needed to form the desired core-shell architecture. Hence, we find that an optimal core-matrix τ_y_ ratio of essentially unity and optimal shell-to-matrix τ_y_ ratio of roughly 10x is required for coaxial embedded printing.

**Figure 2.**
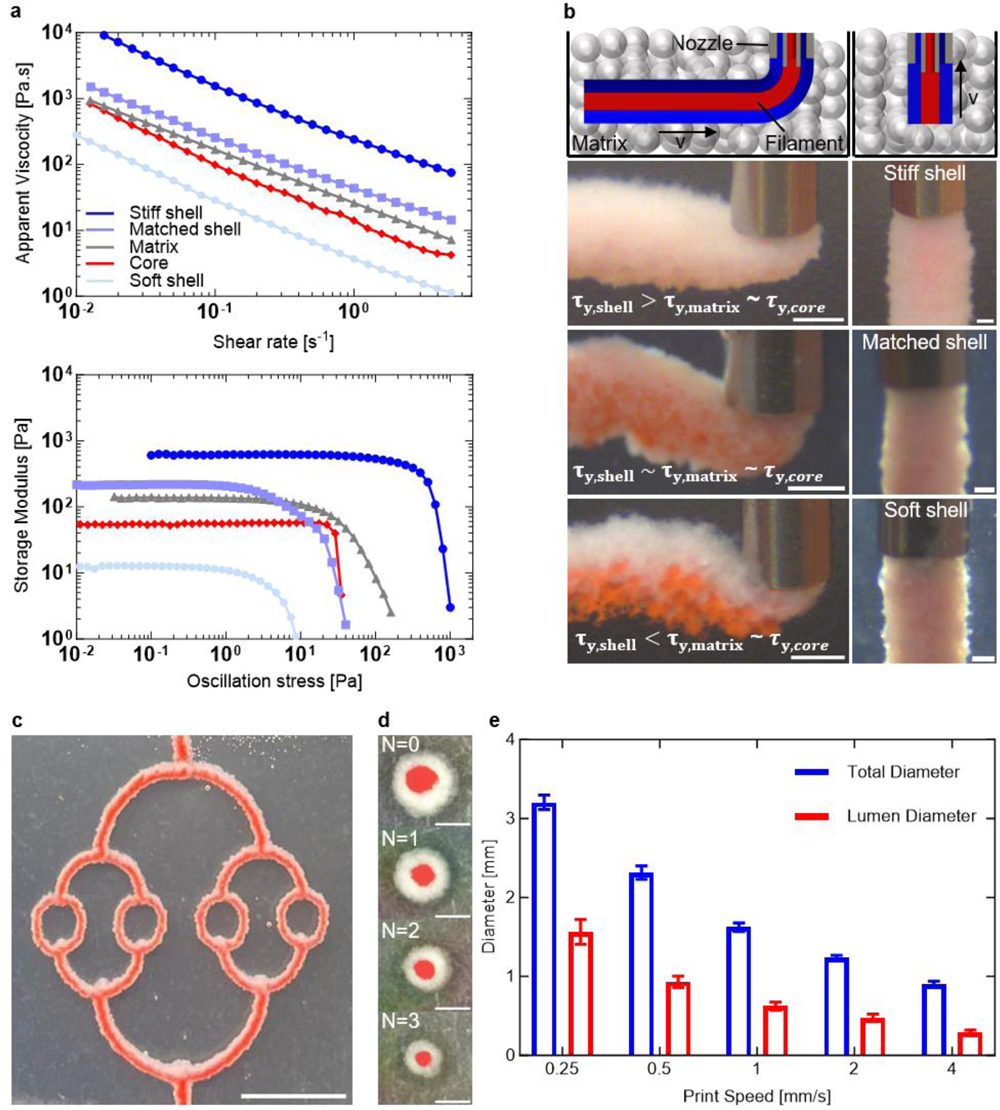
Core, shell, and matrix optimization for coaxial embedded printing. a) Rheological characterization of the core ink, shell inks, and matrix. b) Core-shell filaments printed horizontally (left) and vertically (right) composed of stiff, matched, and soft shell inks within a granular alginate matrix. Scale bars 1 mm (left) and 250 µm (right). c) Optical image of branching vessel network (longitudinally sectioned view) printed into an acellular matrix composed of granular alginate microparticles, in which bifurcating channels follow Murray’s law. Scale bar is 10 mm. d) Optical images (cross-sectional view) of the printed core-shell vessels for each order of the printed network shown in (c). Scale bars are 1 mm. e) Plot of total and core diameters for vessels printed at different speeds at a constant volumetric flow rate.

Next, we explored the effects of key printing parameters by creating a symmetrical 2D vascular network via by co-EMB3DP within our transparent alginate matrix. To emulate native vasculature, we printed coaxial vessels of varying diameter with three generations of branching features (Figure 2c) that obey Murray’s law:^35^

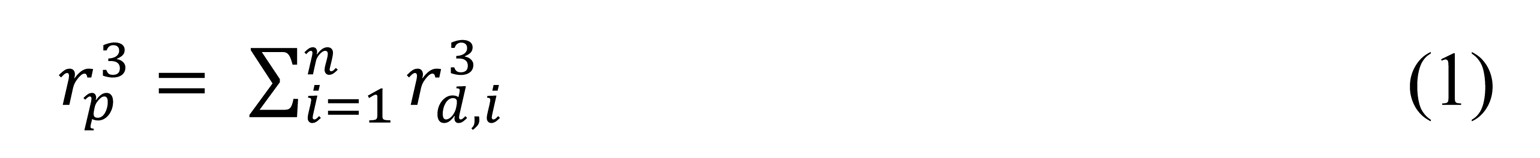

where *r_p_* = radius of the parent vessel and *r_d_* = radius of the daughter vessels branching from the parent vessel. Cross-sectional images of the printed vessels (Figure 2d) reveal that they retain their concentric core-shell architecture across each generation. To produce vessels with total diameters ranging from larger than 3 mm to smaller than 1 mm, we varied the printing speed from 0.25 mm sec^-1^ to 4 mm sec^-1^, while extruding the core and shell inks at a constant volumetric flow rate (Figure 2e). Concomitantly, the core (luminal) diameter decreased from 1.57 mm to 0.29 mm, respectively, over these printing conditions. Alternatively, at a constant printing speed, one can vary the core-to-shell ratio by changing the relative volumetric flow rates of each ink to produce filamentary features ranging from those containing a core-to-shell ink ratio ranging from 0 to 1 (Figure S4).

### Embedding biomimetic vascular networks in µPOROS matrices

To further demonstrate co-EMB3D printing, we designed, printed, and perfused a 3D hierarchical, branching vascular network embedded within an extracellular matrix composed of µPOROS collagen. This matrix is produced by suspending sacrificial gelatin-chitosan microparticles in a pre-polymer collagen solution followed by jamming to induce the desired shear thinning response when locally yielded at an applied shear stress (τ) that exceeds τy ∼ 10 Pa (Figure S5). The µPOROS support matrix and collagen shell ink are held below their gelation temperature for the duration of printing by pumping ice-cold water through a cooling system in the print gasket. The embedded vessels consist of a hierarchically branching network that is patterned in three dimensions and conforms to Murray’s law (Figure 3a-c). Upon printing, the tissue construct is warmed to 37 °C to facilitate collagen gelation and crosslinking in both the shell ink and µPOROS matrix, while simultaneously melting the sacrificial gelatin core (dyed red) (Figure 3b). The vascularized tissue construct is then perfused with PBS (dyed blue) to visualize the perfused, interconnected luminal network (Figure 3c).

**Figure 3.**
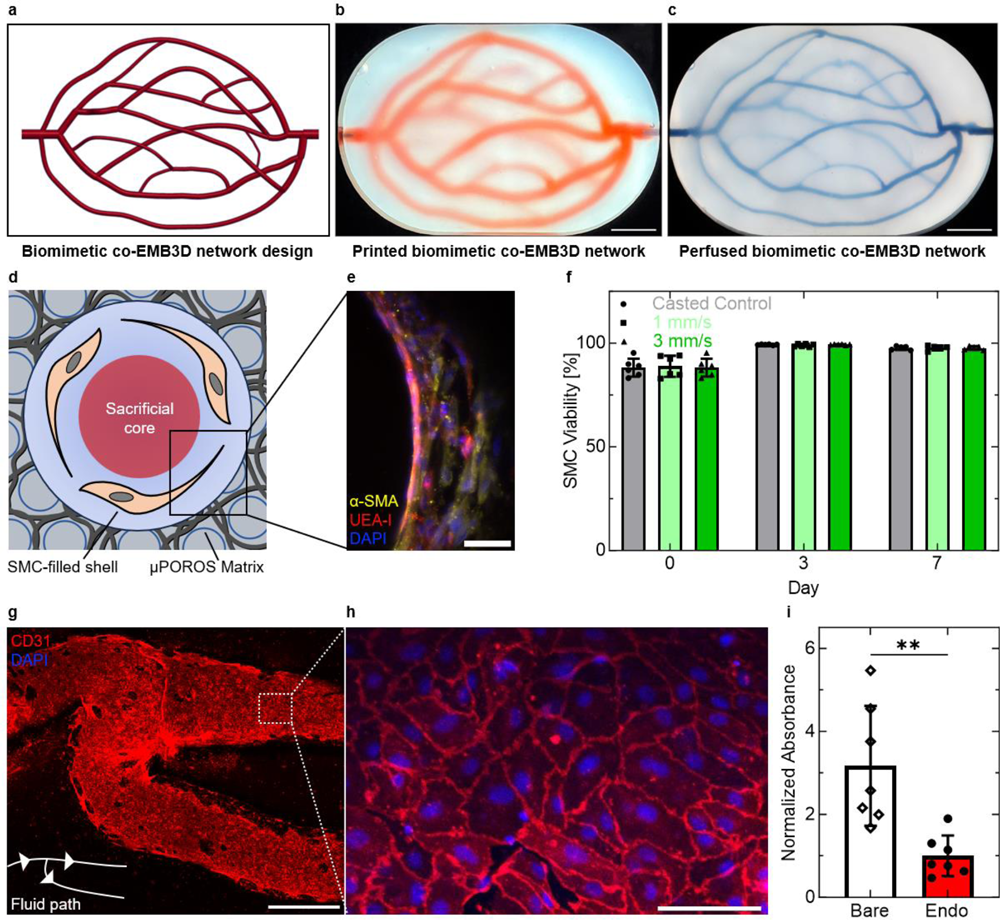
Endothelialization of printed biomimetic blood vessels. a) Rendering of the biomimetic vascular network. b) Printed biomimetic vascular network composed of branching, hierarchical core-shell vessels. c) Perfused biomimetic vascular network after removal of the sacrificial core ink to generate interconnected lumens. Scale bars (b-c) are 10 mm. d) Schematic representation of the as-printed, vessel into the granular µPOROS matrix prior to lumen formation via sacrificial ink removal. e) Cross-section of printed and endothelialized vessel on day 7 of perfusion. Scale bar is 50 µm. f) Live/dead assay of printed smooth muscle cells and as-cast control over 7-day period of vessel perfusion. g) Confocal image (longitudinal cross-section) of a branched, endothelialized vessel network produced by co-EMB3DP. Scale bar is 1 mm. h) Higher magnification, confocal image of confluent endothelium lining the printed and perfused branching vessels. Scale bar is 50 µm. i) Permeability assay of bare and endothelialized vessels following one week of culture.

To further enhance the physiological relevance, we printed a biomimetic vascular network composed of a SMC-laden shell ink surrounding a sacrificial gelatin core ink within this µPOROS matrix (Figure 3d). Upon heating to 37 °C, the SMC-laden shell ink gels to create the blood vessel walls, which surround the interconnected luminal network that forms upon core ink removal. The luminal surfaces are coated with 1% v/v Matrigel on day 2 of perfusion prior to seeding the vessels with endothelial cells. After day 7 of perfusion, the smooth muscle cells remain viable, spread, and wrap around the vessel walls circumferentially akin to the morphology found in the native medial layer (Figure 3e-f, Figure S6).^[18]^ The endothelial cells are arranged in a confluent monolayer with adherent junctions (Figure 3g-h). We then carried out a Miles assay to assess their barrier function.^[36]^ We observed a three-fold decrease in dye diffusion from blood vessels that possess a confluent endothelium compared to the bare (control) vessels (Figure 3i).

### Embedding biomimetic vascular networks in functional cardiac tissues

As a final demonstration, we generated bulk cardiac tissues with biomimetic vasculature via co-SWIFT. We first created hundreds of thousands of cardiac organ building blocks (OBBs) composed primarily of hiPSC-derived cardiomyocytes following previously reported protocols.^[3,37]^ Next, we suspended these cardiac OBBs in a fibrin solution that exhibits a fluid-like response under ambient conditions. Next, we used centrifugation to jam the cardiac OBB solution into a densely cellular matrix (τy ∼ 10 Pa, cell density of ∼200x10^6^ cells mL^-1^) (Figure 4a).^[3]^ We then patterned biomimetic vessels within this cardiac tissue matrix via co-SWIFT printing of the SMC-shell/gelatin sacrificial core inks. Thrombin which is added into the shell ink rapidly gels the fibrin solution surrounding the cardiac OBBs after printing. Upon warming the bulk cardiac tissue to 37°C, the sacrificial gelatin ink which forms the core melts, allowing the seamless removal of the core ink to form an interconnect luminal network. After one day of perfusion, we carried out a live-dead assay on these co-SWIFT cardiac tissues (overall diameter = 2.8 mm and height = 1 cm), which revealed their high cell viability throughout their cross-section. (Figure 4b). On day 2 of perfusion, we seeded endothelial cells onto the luminal surface of the embedded vessels. After day 7 of perfusion, the embedded vessels consist of a confluent layer of endothelial cells surrounded by smooth muscle cells (Figure 4c). These densely cellular, vascularized cardiac tissues begin to contract on the first day of perfusion, while their contractile response increases by roughly three-fold from day 1 to day 5 of perfusion (Figure 4d). Importantly, these co-SWIFT cardiac tissues also exhibit a cardio-effective drug response. Upon perfusion of oxygenated media supplemented with isoproterenol at a concentration of 10 µM, we observed their spontaneous beat frequency doubles.^[38]^ By contrast, the perfusion of media that contains 10 µM blebbistatin arrests beating of these cardiac tissues (Figure 4e-f).^[39]^ To highlight co-SWIFT’s promise for personalized biomanufacturing, we printed a scaled model of the main branches for a patient-specific, left coronary artery (LCA) model. To aid visualization, we simultaneously printed the initial branch and full arterial structure into both our transparent alginate matrix and densely cellular, cardiac OBB matrix (Figure 4g-h, Movie S4). In the future, we plan to generate self-assembled microvascular networks (capillaries) within these co-SWIFT cardiac tissues and promote their anastomosis to printed vessels in vitro to better recapitulate the native myocardium and enhance function.

**Figure 4.**
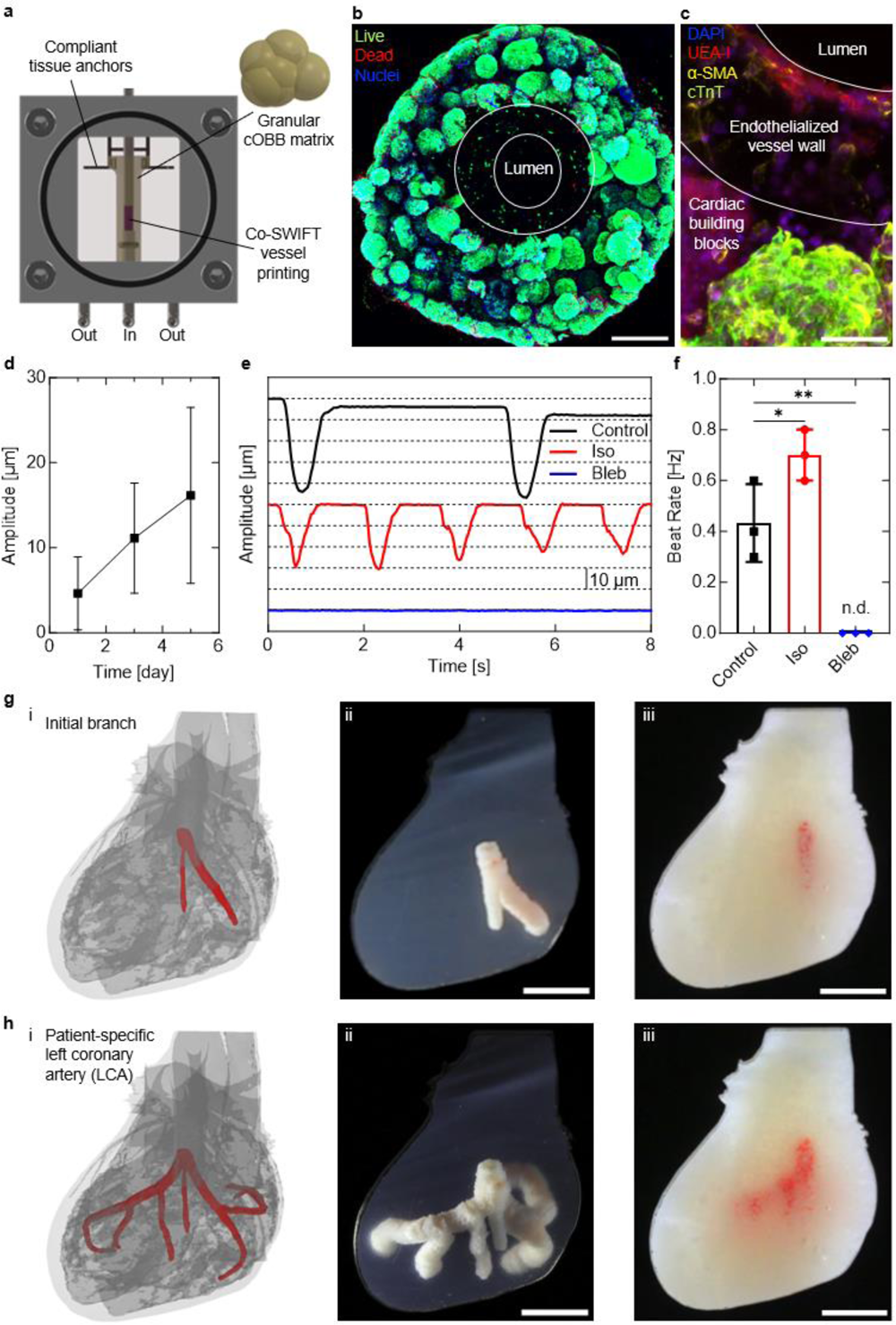
Vascularized cardiac tissues fabricated via co-SWIFT. a) Customized printing and perfusion chamber for cardiac co-SWIFT tissues. b) Live/dead assay of cardiac co-SWIFT tissue. White lines denote the edges of the vessel wall. Scale bar is 500 µm. c) Cross-section of endothelialized cardiac co-SWIFT tissue following five days of culture. Scale bar is 100 µm. d) Beat amplitude of cardia co-SWIFT tissues over time. e) Beat amplitude traces of cardiac co-SWIFT tissues following drug treatment. f) Quantification of spontaneous beating rate of cardiac co-SWIFT tissues following drug treatment. g-h) (i) CAD rendering of printed LCA structure. Printed LCA in alginate microparticles (ii) and in cOBBs (iii) of the initial branch (g) and the completed patient specific LCA (h). Scale bars (g-h) are 5 mm.

## 3. Conclusion

In summary, we have established coaxial-based printing methods for embedding biomimetic vascular networks into both acellular and densely cellular tissue matrices. To demonstrate broad applicability, we tailored the rheological properties of our core, shell, and matrix materials for co-EMB3DP in a granular alginate matrix and microporogen-structured collagen as well as co-SWIFT printing in functional cardiac tissues. Through the design, fabrication, and implementation of customized extended core-shell nozzles, we demonstrated that hierarchical branching vessels composed of smooth muscle cell-laden shell ink that surround a sacrificial core ink could be produced. Such networks possess interconnected lumens (upon removing their sacrificial core), which were wrapped by smooth muscle cells and seeded with endothelial cells to form a confluent endothelium that provides good barrier function. Finally, we created thick cardiac tissues with embedded biomimetic vessels, whose design is guided by patient-specific data. Our work provides an enabling advance for embedding biomimetic vascular networks within soft and living tissue constructs.

## 4. Experimental Section

### Core-shell nozzle design and fabrication

The extended core-shell bioprinting nozzle was designed in Solidworks (Dassault Systemes) and printed on the EnvisionTec D4K printer using HTM140 resin (Desktop Metal). The nozzles were cleaned by connecting a syringe to the luer-lock and purging the fluid paths with 2-propanol (Sigma-Aldrich). A core nozzle (inner diameter = 0.25 mm, outer diameter = 0.52 mm, and length = 3.15 cm) was mated with a shell nozzle (inner diameter = 0.84 mm, outer diameter = 1.27 mm, and length = 1.9 cm) (Nordson EFD). The core nozzle was secured by injecting superglue (Loctite), while the shell nozzle was affixed using epoxy (Loctite).

### Core-Shell Inks

Sacrificial gelatin used for the core and shell inks was prepared by dissolving 300 g Bloom type A gelatin (Sigma-Aldrich) at 15% w/v and stirred at 85 °C for 12 h. This gelatin stock was then adjusted to pH 7.4 using 1 N sodium hydroxide (Sigma-Aldrich). The gelatin was then sterile filtered and stored at 4 °C for up to 6 months. To prepare the core and shell inks, stock 15% w/v gelatin and 70 mg mL^-1^ neutral collagen (LifeInk220, Advanced Biomatrix) respectively were diluted with different amounts of phosphate buffered saline with calcium and magnesium (PBS) (Corning) to the final concentrations in **Table 1**. To provide visual contrast between the core and the shell inks, either red pigment (Gamblin) or red food coloring (Ward’s Science) was added to the core inks.

**Table 1.**
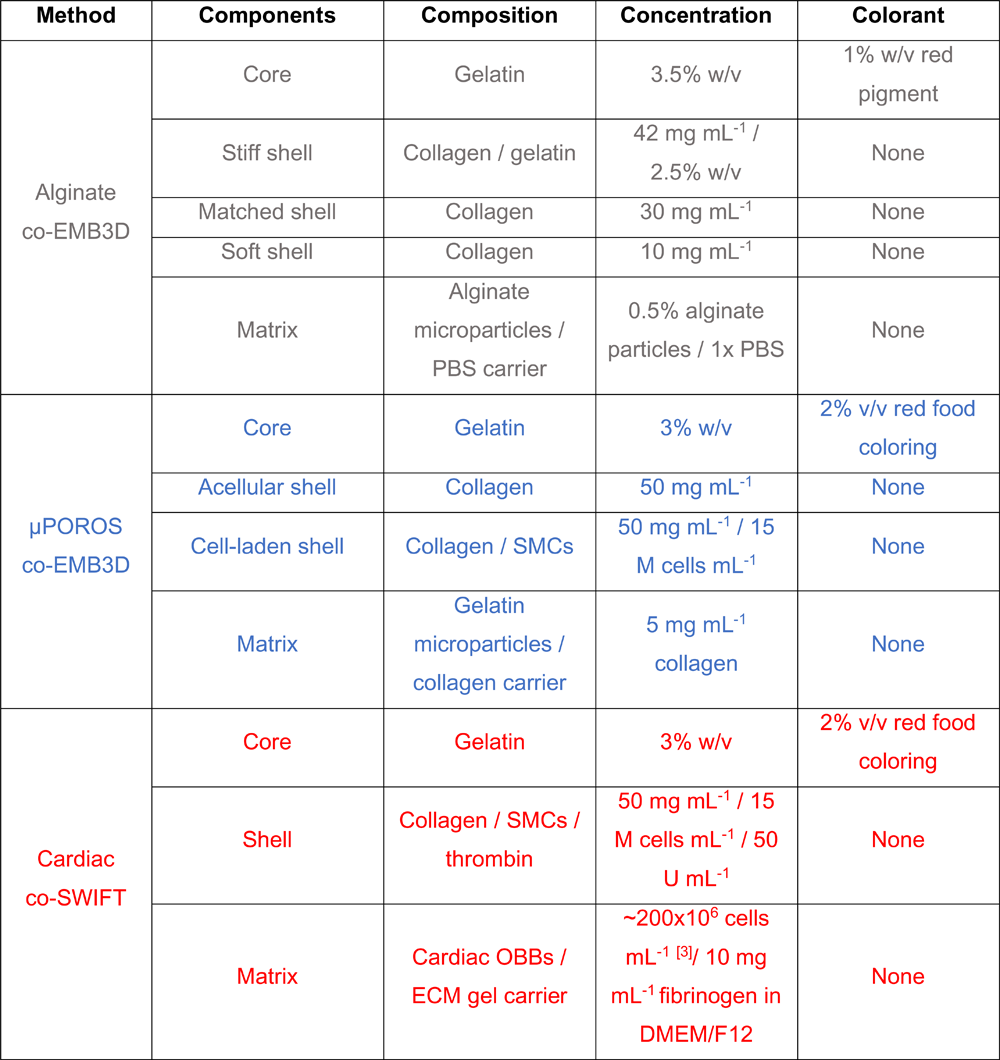
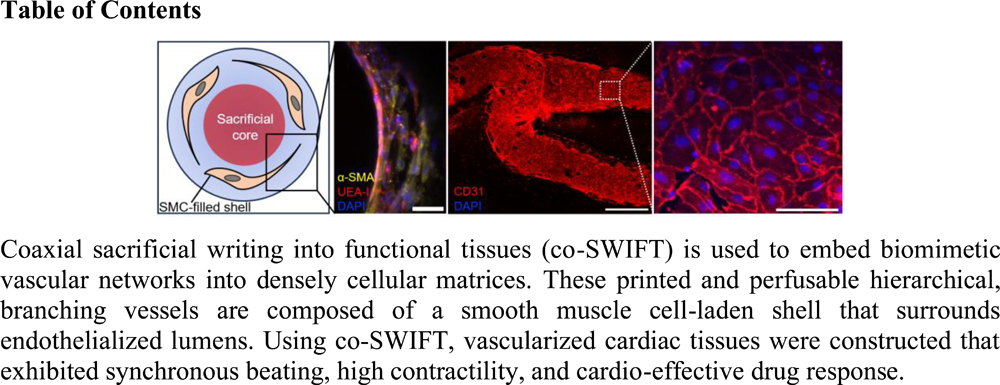
Composition of co-SWIFT Materials.

### Granular Alginate Matrices

Alginate solutions were generated by dissolving medium viscosity alginic acid sodium salt (Sigma-Aldrich) in deionized water. Granular alginate particles (diameter = 190 µm ± 19.2 μm) were fabricated by injecting 0.5% alginate or 2% alginate solution from a lavender 45° nozzle (Nordson EFD) at a flow rate of 300 µL min^-1^ into a 2 psi air stream controlled by a pressure box (Nordson EFD) through a purple 0.5 in nozzle (Nordson EFD). The alginate droplets were deposited into a gelation bath containing 100 mM CaCl_2_ (Sigma-Aldrich) and 5% ethanol (KOPTEC), where they were then crosslinked for 1-3 h prior to being washed and stored in an aqueous salt solution containing 2 mM CaCl_2_. These granular alginate particles were stored at 4 °C for up to 3 months before use.

To create printable matrices, the granular alginate particles were first swelled in PBS for 90 min and then centrifuged at 30g for 3 min. The supernatant was removed and the jammed particles were mixed with a serological pipette prior to loading them into the printing chamber or onto a controlled-shear rheometer. To quantify the volume fraction of alginate particles within these printable (jammed) matrices, 0.05% w/v 2 MDa TRITC-dextran (Thermo-Fisher) was added to the alginate solution prior to its consolidation. Confocal microscopy (Zeiss) coupled with image analysis was used to determine the volume fraction of granular alginate particles within the printable matrices. In addition, videos were generated from z-stack confocal images using a custom MATLAB script (MathWorks).

### µPOROS matrices

Our µPOROS matrix consists of sacrificial gelatin microparticles and prepolymer collagen I. Following our published protocol^[33]^, we generated sacrificial microparticles by dissolving 2% w/v gelatin type A (Sigma-Aldrich), 0.25% w/v Pluronic F-127 (Sigma-Aldrich), and 0.1% w/v chitosan (Sigma-Aldrich) in 51% v/v ethanol (Sigma-Aldrich) while stirring at 45 °C. The pH was adjusted to 6.32 using 1 N NaOH. The sacrificial gelatin microparticles were removed from heat and stirred overnight. The next day, the microparticles were homogenized mechanically and washed 3x in PBS. The microparticles (∼30-50 μm in diameter) were stored at 4 °C for up to 4 months before use. Immediately prior to co-SWIFT printing, the sacrificial gelatin microparticles were centrifuged at 2000g for 3 min and the supernatant was removed. The particles were resuspended in 5 mg mL^-1^ type I collagen (Advanced Biomatrix), transferred to 10 mL syringes, and centrifuged at 3000g for 5 min at 4 °C. The supernatant was removed and the µPOROS matrix was passed between two syringes using a syringe coupler 20 times to homogenize the matrix. The µPOROS matrix was stored in ice-water until it was used for printing or rheological characterization.

### Rheological characterization

All rheological measurements were carried out on a controlled stress-controlled rheometer (DHR-3, TA Instruments) with a 25 mm diameter aluminum parallel plate geometry with 60 grit sandpaper attached to the surface to prevent slipping. Gap heights of 250 µm, 1 mm, and 2 mm, and were used for the inks, µPOROS matrix, and granular alginate matrix, respectively. Shear and oscillatory measurements for both inks and the µPOROS matrix were carried out at 2 °C, while measurements on the granular alginate matrix were performed at 20 °C. Apparent viscosity curves were collected by performing flow sweeps at shear rates ranging from 10 s^-1^ to 0.001 s^-1^, while oscillatory measurements were performed at 0.5 Hz from 0.005 Pa until yielding.

### Primary cell culture

Primary human umbilical vein endothelial cells (Lonza) and aortic smooth muscle cells (Cell Systems) were cultured in endothelial growth medium (EGM-2, Lonza) and VascuLife smooth muscle cell medium (LifeLine Cell Technology) respectively. Medium was refreshed every other day until the cells were 80% confluent. The cells were passaged by first rinsing with PBS without calcium and magnesium (PBS-/-) (Corning), then adding one quarter culture volume of 0.05% trypsin/EDTA (Gibco) to the flask for 4 min at 37 °C. The 0.05% trypsin/EDTA was quenched using DMEM/F12 with 10% fetal bovine serum (FBS) (Gibco). The cells were then centrifuged at 220g for 3 min. The supernatant was removed and the cells were split into preprepared flasks at a ratio ranging from 1:3 to 1:5. All primary cells were used from passage 4 to passage 7.

### Embryoid body formation

BJFF iPSCs (provided by S. Jain at Washington University) were cultured on stem-cell qualified growth factor reduced Matrigel (Corning) in mTeSR Plus stem cell medium (STEMCELL Technologies) in a 37 °C/5% CO_2_ incubator. Once colonies reached 70-80% confluency, they were rinsed once in PBS-/-. ReLeSR (STEMCELL Technologies) was added to the flask and immediately aspirated away. The cells were transferred to the incubator for 7 min, before they were gently rinsed with culture medium and added to a freshly prepared flask at a ratio of 1:8. The iPSCs were lifted from the flask to form embryoid bodies (EBs) using the same method as for passaging, but were seeded in mTeSR Plus medium supplemented with 10 µM Y27632 (BioGems) into a non-adherent T25 flask (Corning) at a ratio of 112.5 cm^2^ adherent culture area per non-adherent T25 flask on day -5 of differentiation. The flasks were then placed on an orbital shaker at 55 RPM. Medium was changed the next day with fresh mTeSR Plus without Y27632, then every other day until day 0.

### Cardiac building blocks

A modified protocol^3^ from Lian et al.^37^ was used to differentiate the EBs into cardiac spheroids. On day 0, differentiation was initiated by adding cardiac differentiation medium (CDM) composed of RPMI 1640 (Gibco) and 2% B27 without insulin (Gibco) and supplemented with 5 µM CHIR99021 (BioGems). The same medium was refreshed on day 1. On day 2 of differentiation, CDM with CHIR99021 was removed and replaced with CDM. On days 3 and 4, CDM with 2 µM iWR1 (BioGems) was added. On day 5, the cells were cultured in CDM until beating was observed (day 6 or 7), after which CDM was replaced with cardiac maturation medium (CMM) composed of RPMI 1640 and 2% B27 with insulin (Gibco) and refreshed daily until the cardiac spheroids were used for co-SWIFT experiments (day 10-12).

### Print path generation

Complex print paths were first designed in Solidworks and then exported to MATLAB. To generate the print path for the patient derived left coronary artery geometry (https://3d.nih.gov/entries/3dpx-012589, Karolina Stepniak, 2019), the structure was first downloaded from the NIH 3D print exchange and imported into Solidworks. A custom MATLAB script was used to translate the point data into G-code with the desired flow rates and print parameters, and the geometry scaled as needed. All other print paths were generated directly in G-code. Each print path was imported to A3200 motion control software (Aerotech) used to control our customized, multi-material 3D bioprinter.

### Printing and perfusion chambers

To facilitate coaxial printing and perfusion of embedded vasculature, customized chambers were either machined from polycarbonate (McMaster-Carr) or printed via stereolithography using BioMed Clear resin (Formlabs). In both cases, a watertight seal was formed using O-rings (McMaster-Carr), which were compressed using laser-cut acrylic plates (McMaster-Carr). The metal inlet and outlet pins (Nordson EFD) were epoxied (Loctite) to the main body of the culture chamber. The compliant spring arms were printed using EnvisionTec D4K printer with a HTM140 resin. Before sterilization, the compliant support arms were inserted into the gasket. The culture chambers and compliant support arms were autoclaved, while the acrylic windows were sterilized in 70% ethanol for a minimum of 30 min before use.

### co-SWIFT printing

The night before printing, 300 g Bloom type A gelatin which was dissolved at 15% w/v 70 °C for 1 h was diluted to 5% w/v using DMEM/F12 with HEPES (Gibco) and supplemented with CaCl_2_ to a final concentration of 2.5 mM and 5 U mL^-1^ thrombin. A 3D printed mold in the desired shape of the tissue was coated with 10% Pluronic F-127 (Sigma-Aldrich) and inserted into the gasket. The gelatin-thrombin solution was used to fill the gasket around the mold. The gaskets were then stored at 4 °C overnight. On the day of printing, the molds were removed from the gelatin and the negative cavity was rinsed 3x with PBS. The chambers were then held at 4 °C.

Immediately prior to printing, an anchor gel which was used to affix the co-SWIFT tissue to the perfusion pin was prepared from two precursor solutions to prevent premature polymerization. Part 1 of the precursor solution contained 20 mg mL^-1^ fibrinogen (Merck) diluted in DMEM/F12 with HEPES. Part 2 of the precursor solution consisted of 2.5 mM CaCl_2_, 0.5 U mL^-1^ thrombin, and 20 mg mL^-1^ transglutaminase (Moo Gloo TI). Part 1 and part 2 were mixed in equal volume and allowed to polymerize at the base of the inlet pin. The extracellular matrix (ECM) gel, which provided immediate structural support for the co-SWIFT tissue upon crosslinking after printing, consisted of 10 mg mL^-1^ fibrinogen and 2.5 mM CaCl_2_ (Sigma-Aldrich) diluted in DMEM/F12 with HEPES.

Once the anchor gel was added to the chamber, the cOBBs were rinsed with 3:1 v/v ECM gel, centrifuged at 30g, and the supernatant was aspirated. The cOBBs were resuspended in 1:1 v/v ECM gel to cOBBs and transferred to a 1 mL disposable syringe (BD Biosciences). The cOBBs were centrifuged at 100g for 3 min and the supernatant removed. The resulting jammed cOBBs were then dispensed into the mold using an olive nozzle (inner diameter = 1.54 mm, Nordson EFD). Embedded vascular networks were rapidly printed within cOBB matrices via co-SWIFT of the sacrificial core ink and matched shell ink filled with SMCs. The customized printing and perfusion chambers were then transferred to the incubator at 37 °C to promote rapid polymerization of the collagen and fibrin within the shell ink and ECM gel, respectively, while the sacrificial gelatin ink in the core and the surrounding chamber liquify. After 20 min, the co-SWIFT tissues were connected to a peristaltic pump (MasterFlex), and the co-SWIFT medium consisting of equal volumes of CMM and vessel co-culture medium with 1:250 aprotinin (EMD Millipore) and 1x Antibiotic-Antimycotic (Gibco) was used to evacuate the sacrificial gelatin from the vessel and chamber at a flow rate of 100 µL min^-1^. Once the sacrificial gelatin was removed, the flow rate was slowly increased to 250 µL min^-1^ until day 2 when the tissues were endothelialized (as described below), and the flow rate was increased to 500 µL min^-1^ for the duration of culture. The co-SWIFT medium was refreshed every other day. ***Endothelialization of co-SWIFT vessels***: Vessels were coated with a 1% v/v Matrigel solution in either vessel co-culture medium or co-SWIFT culture medium for 2 h before endothelialization. HUVECs were lifted from the flask as previously described, then injected into the vessel at 20x10^6^ cells mL^-1^. These endothelial cells were allowed to attach for 80 min without flow at 37 °C during which the culture chamber was rotated 90° every 10 min to ensure even coating of the luminal surface. Flow was resumed at 50 µL min^-1^ for 10 min, then slowly ramped up to its steady-state value of 500 µL min^-1^ over a 20 min period.

### Immunofluorescent Staining and Confocal Imaging

co-SWIFT vessels and cardiac co-SWIFT tissues were fixed in 4% paraformaldehyde (PFA) (Electron Microscopy Sciences) for 30 min or 45 min, respectively. Tissues were washed 3x for a minimum of 15 min in PBS before immunofluorescent staining. Permeabilization and blocking were performed for two hours in PBS containing 0.125% Triton X (Sigma-Aldrich), 0.5% bovine serum albumin (BSA) (Miltenyi Biotech), and 2% donkey serum (Sigma-Aldrich). Primary antibodies [cTnT (ab45932), αSMA (ab7817), CD31(ab9498) (Abcam)] were added at 1:200 in PBS with 0.125% Triton X and 0.5% BSA at 4 °C for 12 to 24 h. The constructs were then washed 3x for a minimum of 15 min in PBS before secondary antibody incubation. Alexa Fluor Plus conjugated secondary antibodies (Invitrogen) and UEA-I conjugated with fluorescein (Vector Laboratories) were then added in PBS with 2% donkey serum either for 2 hours at room temperature or at 4 °C overnight. 4’,6-diamidino-2-phenylindole (DAPI) (Thermo Fisher) was added for 30 min at room temperature before the secondary antibodies were washed out in PBS 3x for 15 min. Constructs were imaged on an upright confocal microscope (Zeiss).

### Cell viability assays

The viability of smooth muscle cells encapsulated in the shell ink was assessed by first removing the cell culture medium from the customized printing and perfusion chamber and then adding PBS with ethidium homodimer and calcein AM at 1x working concentrations of 0.5 µL mL^-1^ and 2 µL mL^-1^, respectively, based on manufacturer recommendations (Invitrogen). The tissues were incubated at 37 °C for 30 min before imaging on an upright confocal microscope using a 10x water immersion objective. Quantitative image analysis was performed using Imaris (Oxford Instruments). To quantify cell viability, the cardiac co-SWIFT tissues were removed from flow after 24 h and sectioned into cylinders (roughly 1 mm in height and 2.5 mm in diameter) in a chamber containing ice-cold, co-SWIFT culture medium. Next, 50% of this medium replaced with an equal volume of ethidium homodimer and calcein AM at a 2x working concentration. Hoechst solution (Invitrogen) was added at a final concentration of 0.25 µL mL^-1^. The tissues were incubated at 37 °C for 30 min before imaging on a confocal microscope with a 5x non-immersion objective.

### Barrier function assay

A Miles permeability assay was performed to assess barrier integrity of the endothelial monolayer on the surface of the co-SWIFT vessels. After 1 week of culture, a 1% w/v solution of Evan’s blue dye (Chem-Impex International) was dissolved in PBS. It was diluted 1:9 in vessel co-culture medium (final dye solution). The final dye solution was perfused through the vessel for 20 min at a flow rate of 500 µL min^-1^ before the vessel was flushed with PBS for 5 min at the same flow rate to remove excess dye from the lumen of the vessel. The vessel construct was removed from the culture chamber and weighed on an analytical balance. The construct was then dissolved in 200 µL formamide (G-Biosciences) for 48 h at room temperature to recover the dye. The absorbance at 630 nm was recorded on the SynergyHT plate reader (BioTek) and the values were normalized to the weight of the construct.

### Cardio-effective Drug Response

On day 10 of culture, either isoproterenol (Sigma-Aldrich) or blebbistatin (Sigma-Aldrich) was delivered intraluminally to the co-SWIFT cardiac tissues at a concentration of 10 µM for 30 min. After 30 min, videos of the cardiac co-SWIFT tissues were collected on a VHX-2000 digital microscope (Keyence). The resultant videos were analyzed using the open-source software, Tracker (https://physlets.org/tracker/).

## Acknowledgements

The authors gratefully acknowledge support from the Vannevar Bush Faculty Fellowship Program sponsored by the Basic Research Office of the Assistant Secretary of Defense for Research and Engineering through the Office of Naval Research Grant N00014-21-1-2958 and the National Science Foundation through CELL-MET ERC (#EEC-1647837). The authors thank P. Lustenberg for early contributions to the coaxial nozzle design, A. Lu, J. Wilt, and J. Ahrens for useful discussions, and L.K. Sanders for videography.

## Supporting Information

**Figure S1.**
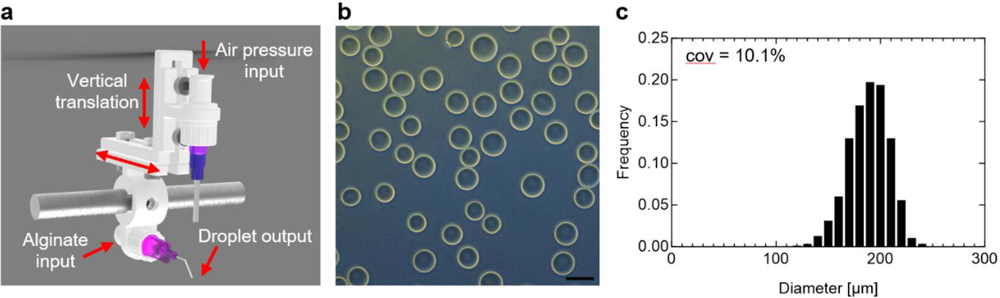
Granular alginate production. a) Schematic representation of alginate particle fabrication jig. b) Representative phase contrast image of the granular alginate particles. Scale bar is 250 µm. c) Histogram of alginate particle sizes obtained.

**Figure S2.**
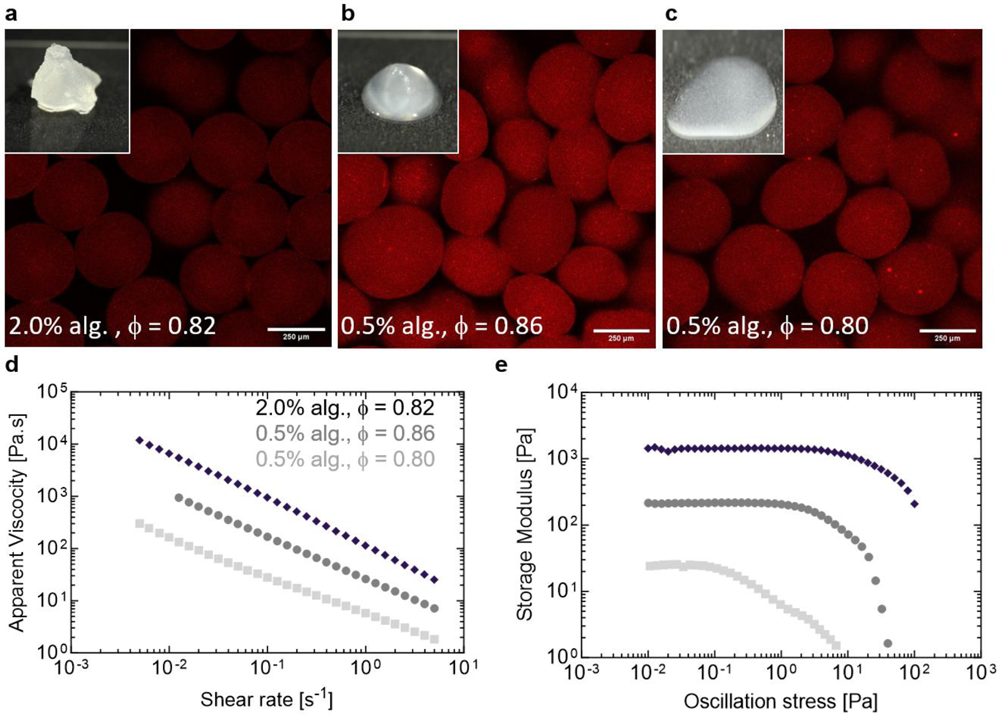
Structure and rheological behavior of granular alginate matrices. a-c) Representative confocal images of a 3D reconstruction of fluorescently dyed alginate particles within matrices of varying composition. Insets show the corresponding optical images of granular alginate matrices of varying composition. d) Log-log plot of the apparent viscosity as a function of shear rate for these granular alginate matrices. (e) Log-log plot of the storage modulus as a function of oscillatory shear stress for these granular alginate matrices.

**Figure S3.**
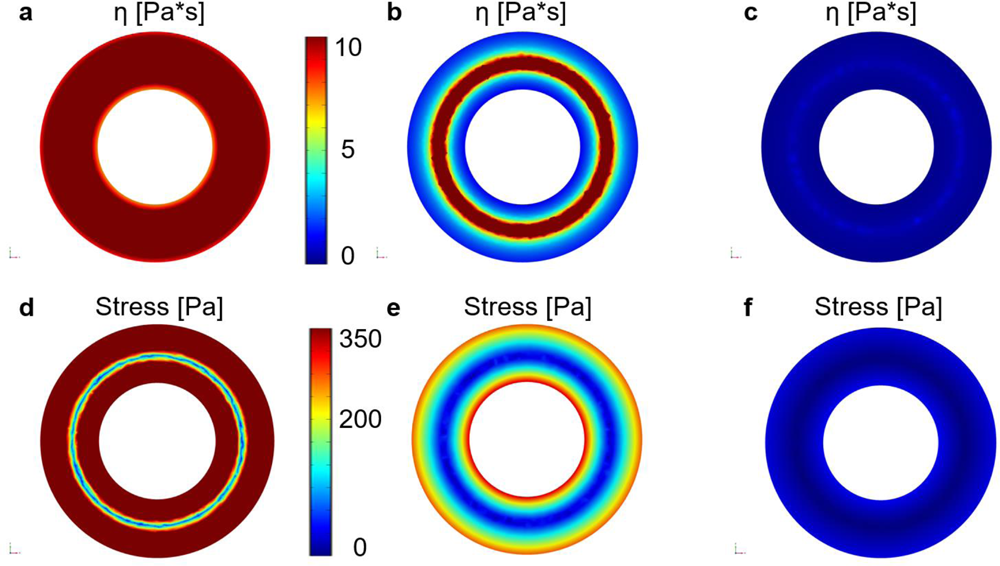
Modeling shell ink flow within coaxial printheads. a-c) From left to right: COMSOL simulations of viscosity as a function of radial position within the coaxial nozzles for the stiff, matched, and soft shell inks (see Figures 1). d-e) From left to right: COMSOL simulations of shear stress as a function of radial position within the coaxial nozzles for the stiff, matched, and soft shell inks. Shell thickness is 160 µm.

**Figure S4.**
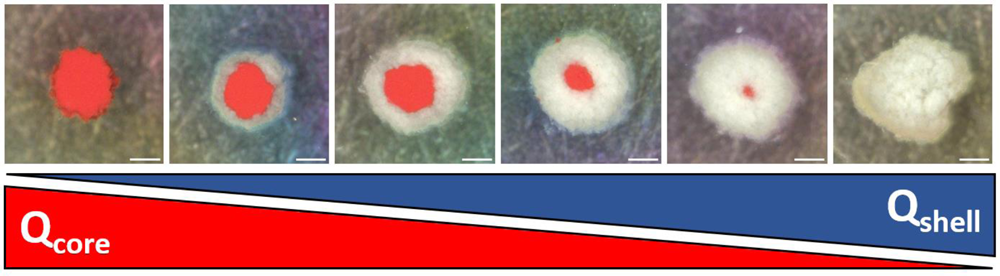
Controlling biomimetic vessel architecture via co-SWIFT. Optical images of vessel cross-section ranging from 100% core ink (left, red) to 100% shell ink (right, white) produced by varying their respective volumetric flow rates at a constant printing speed of 1 mm s^-1^. Scale bars are 250 µm.

**Figure S5.**
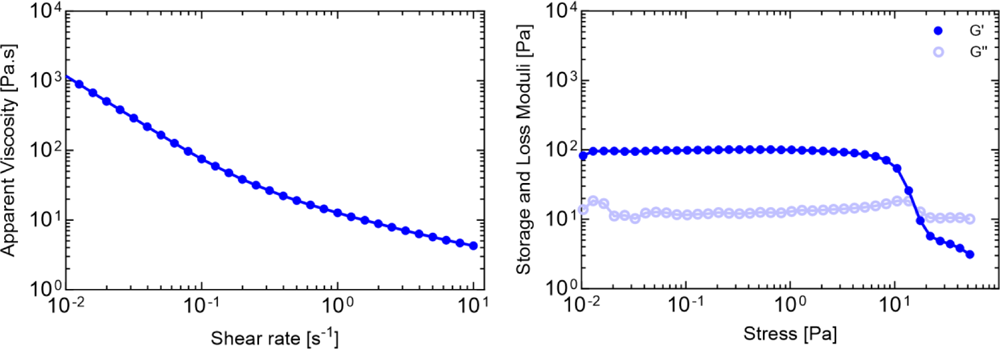
µPOROS matrix rheology. a) Flow sweep of the µPOROS co-SWIFT matrix b) Amplitude sweep of the co-SWIFT matrix

**Figure S6.**
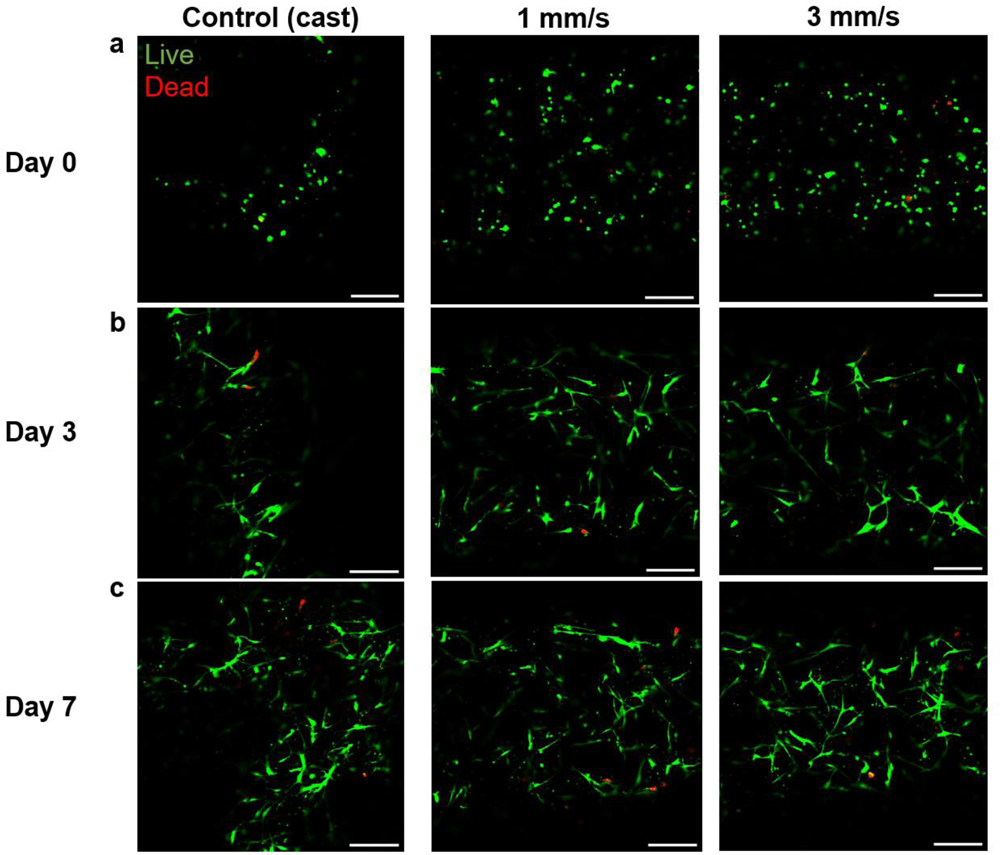
Smooth muscle cell viability in printed vessels. Live/dead assays on (a) day 0, (b) day 3, and (c) day 7 of culture. Scale bars are 250 µm.

**Movie S1.**
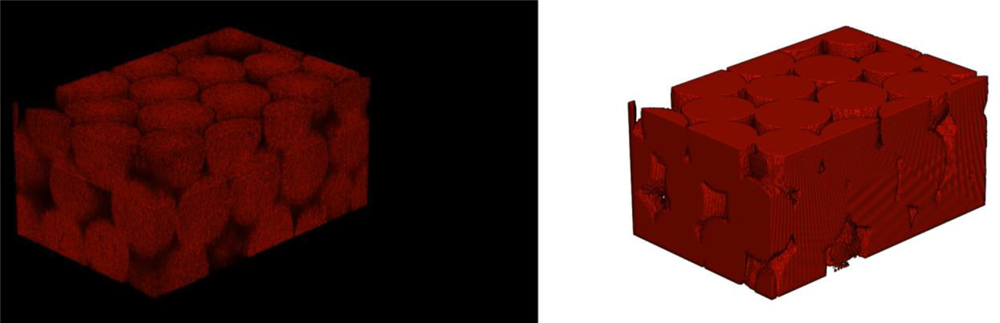
3D structure of the stiffest granular alginate matrix. Video of the confocal image z-stack and MATLAB reconstruction for the 2%, Φ = 0.82 granular alginate matrix loaded with fluorescent dextran.

**Movie S2.**
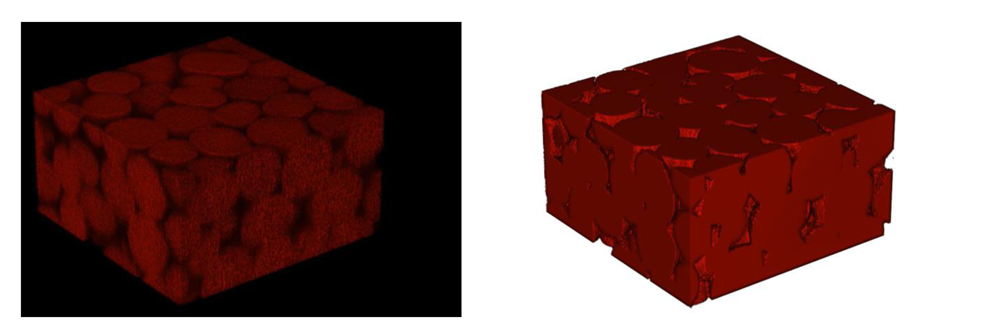
3D structure of the granular alginate matrix with intermediate stiffness. Video of the confocal image z-stack and MATLAB reconstruction for the 0.5%, Φ = 0.86 granular alginate matrix loaded with fluorescent dextran.

**Movie S3.**
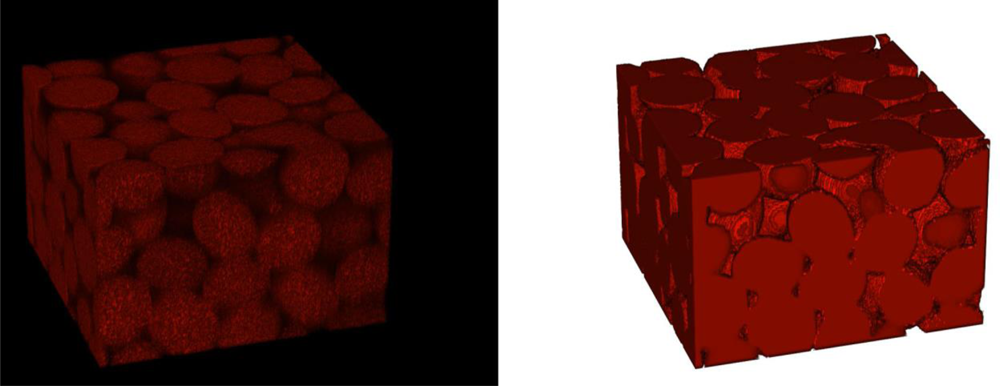
3D structure of the softest granular alginate matrix. Video of the confocal image z-stack and MATLAB reconstruction for the 0.5%, Φ = 0.8 granular alginate matrix loaded with fluorescent dextran.

**Movie S4.**
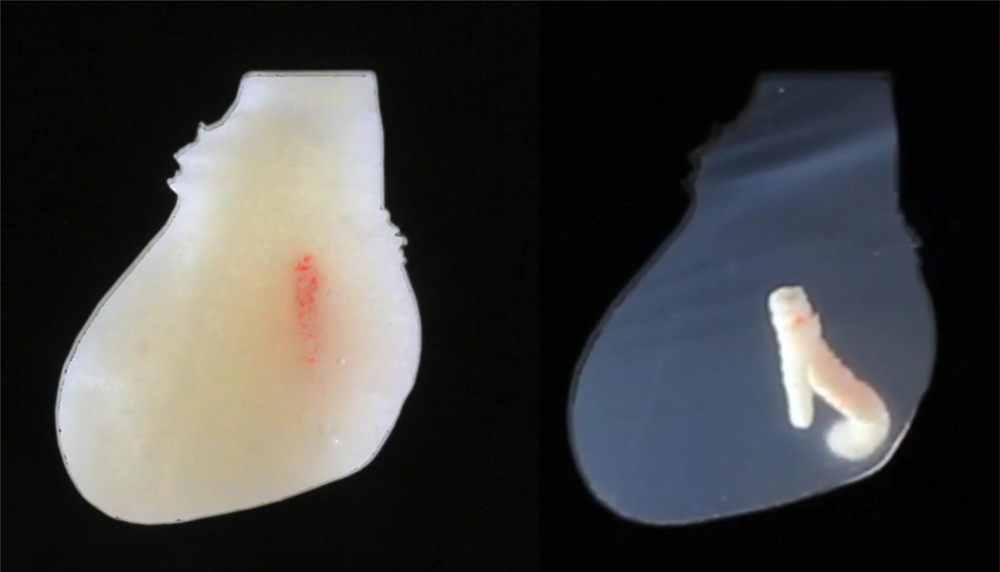
Coaxial embedded printing of a patient-specific LCA model in acellular and living matrices. Side-by-side video of the patient-specific LCA being printed into our functional cardiac tissue (left) and granular alginate matrix (right).

